# The open field assay is influenced by room temperature and by drugs that affect core body temperature

**DOI:** 10.1101/2022.10.17.512128

**Authors:** Jessica A. Jiménez, Eric S. McCoy, David F. Lee, Mark J. Zylka

**Affiliations:** UNC Neuroscience Center, The University of North Carolina at Chapel Hill, Chapel Hill, NC 27599, USA; UNC Curriculum in Toxicology and Environmental Medicine, The University of North Carolina at Chapel Hill, Chapel Hill, NC 27599, USA; Department of Cell Biology & Physiology, The University of North Carolina at Chapel Hill, Chapel Hill, NC 27599, USA

## Abstract

The open field assay is used to study anxiety-related traits and anxiolytic drugs in rodents. This assay entails measuring locomotor activity and time spent in the center of a chamber that is maintained at ambient room temperature. However, the ambient temperature in most laboratories varies daily and seasonally and can differ substantially between buildings. Here, we sought to evaluate how varying ambient temperature and core body temperature (CBT) affected open field locomotor activity and center time of wild-type (WT) and *Trpm8* knock-out (*Trpm8*^-/-^) mice. TRPM8 is an ion channel that detects cool temperatures and is activated by cooling agents, including icilin and menthol. We found that the cooling agent icilin increased CBT and profoundly reduced distance traveled and center time of WT mice relative to vehicle controls. Likewise, cooling the ambient temperature to 4°C reduced distance traveled and center time of WT mice relative to *Trpm8*^-/-^ mice. Conversely, the TRPM8 antagonist (M8-B) reduced CBT and increased distance traveled and center time of WT mice when tested at 4°C. Predictably, the TRPM8 antagonist (M8-B) had no effect on CBT or open field behavior of *Trpm8*^-/-^ mice. The anxiolytic diazepam reduced CBT in both WT and *Trpm8*^-/-^ mice. When tested at 4°C, diazepam increased distance traveled and center time in WT mice but did not alter open field behavior of *Trpm8*^-/-^ mice. Our study shows that environmental temperature and drugs that affect CBT can influence locomotor behavior and center time in the open field assay, highlighting temperature (ambient and core) as sources of environmental and physiologic variability in this commonly used behavioral assay.

## Introduction

Drugs used to treat neurological and neuropsychiatric disorders are typically evaluated in rodent models for safety and efficacy prior to use in humans (1). Characterizing animal models of neuropsychiatric disorders often relies on behavioral traits such as motor function, social interactions, anxiety-like and depressive-like behavior, substance dependence, and various forms of cognitive function (2). Due to the complexity of most behavior tests, researchers must carefully consider the sources of variability introduced by experimenters, testing environments, and intraspecies differences.

Rodent physiology and behavior is influenced by environmental temperature (3, 4). For example, an innocuous cold stimulation at 15°C altered sleeping, rearing, climbing, and eating behavior in wild-type (WT) mice (5). This cold stimulation did not alter these behaviors in mutant mice lacking the Transient Receptor Potential Subfamily M Member 8 (TRPM8) cation channel, which is a receptor for menthol and icilin (mint-derived and synthetic cooling compounds, respectively) and plays an important role in thermosensation (5, 6). Additionally, mice deficient in uncoupling protein 1 (UCP-1), a key metabolic regulator highly expressed in brown adipose tissue, were reported to display selective enhancement of anxiety-related behavior exclusively under thermogenic conditions (23°C), but not at thermoneutrality (29°C) (7).

Environmental temperature sensation and perception is also influenced by core body temperature (CBT). Alterations to CBT can be a consequence of physiological changes associated with disease state, exercise, metabolic function, and hormonal changes. Further, numerous drugs can affect body temperature including barbiturates, cyclic antidepressants, hypoglycemic agents, opioids, antihistamines, and anticholinergic drugs (8–11). It is currently unclear if drugs such as these, which are used to treat neurological and neuropsychiatric disorders, affect CBT directly or indirectly, and if the behavioral tasks that are commonly used to study these drugs are influenced by changes in CBT.

Here we sought to evaluate how ambient temperature and changes in CBT influence locomotor activity and center time in the open field assay—an assay that is commonly used to study anxiolytic drugs and animal models of anxiety. In addition to using WT mice, we also used *Trpm8*^-/-^ mice which lack the primary receptor for cool temperature sensation in mammals, as well as drugs that activate (icilin) or antagonize (M8-B) this receptor. Diazepam was also evaluated as a model anxiolytic drug. By precisely controlling environmental temperature and TRPM8 activity (genetically and pharmacologically), we found that commonly used measures associated with the open field assay are profoundly sensitive to ambient temperature and CBT. To enhance rigor and reproducibility, we recommend that ambient and core temperature be precisely controlled when performing the open field assay. Moreover, drugs that increase activity and center time in the open field test may do so via thermoregulatory mechanisms, independent of effects on anxiety.

## Methods

### Mice

Animal protocols in this study were approved by the Institutional Animal Care and Use Committee at the University of North Carolina at Chapel Hill and were performed in accordance with these guidelines and regulations. All data presented in this study are from mice obtained from crossing *Trpm8*^+/-^ male with *Trpm8*^+/-^ female mice. *Trpm8* mutant mice were obtained from Jackson Laboratories (*B6.129P2-Trpm8^tm1Jul^/J;* stock #008198). Mice were raised in a facility with a 12 h:12 h light:dark cycle with *ad libitum* access to food (Teklad 2020X, Envigo, Huntingdon, UK) and water. Genomic DNA was isolated from tail clips using Proteinase K. Genotyping was performed by polymerase chain reaction (PCR) amplification of genomic DNA with primers: WT Forward 5’-CCT TGG CTG CTG GAT TCA CAC AGC-3’, Mutant Reverse 5’-CAG GCT GAG CGA TGA AAT GCT GAT CTG-3’, WT Reverse 5’-GCT TGC TGG CCC CCA AGG CT-3’.

### Drug administration

Drugs or control (saline or DMSO) were administered intraperitoneally (i.p.). Icilin (I9532-50MG, Sigma) was dissolved in DMSO and administered at 50 mg/kg body weight (bw). Diazepam (RXDIAZEP5-10, Shop Med Vet) was diluted in saline to 1 mg/mL administered at 2 mg/kg bw. M8-B hydrochloride (SML0893-25MG, Sigma) was dissolved in DMSO to 6 mg/ml and administered at 12 mg/kg bw.

### Body temperature

CBT was assessed using a Digi-Sense Thermocouple Meter (Fisher 13-245-293). Mice were acclimated to the procedure 2x each day for one week prior to testing.

Temperature was measured 30, 60 and 90 minutes post drug administration for diazepam and M8-B. To assess the effects of icilin on CBT, measures were taken every 15 minutes.

### Open-field test

Exploratory activity in a novel environment was assessed by a 1 h trial in an open-field chamber (45 cm × 45 cm × 40 cm) 30 minutes post icilin or diazepam administration, and 1 h following M8-B administration. The total distance moved by each mouse in the open arena, and time spent in the center region of the open-field, were recorded by camera (Sony) connected to the EthoVision software (Noldus Wageningen). Testing was performed at room temperature (23°C) or in the cold room (4°C).

## Results

### Room temperature in a laboratory setting must be well-controlled to eliminate daily and seasonal fluctuations

We found that the ambient temperature varied throughout the day (not shown), week, and season in a room that we previously used for behavioral studies (Figure 1A,B, Building 1). Temperature fluctuations are presumably common in laboratory settings because building heating, ventilation, and air conditioning are set to maximize human comfort during the work day and minimize energy use during off-peak hours, like evenings and on weekends. Moreover, room temperature was over 4°C warmer in the winter months and 2°C warmer in the summer months in a different laboratory located in a different building (Building 2; Figure 1B,C). Temperature differences over days, seasons, and buildings represent a major source of variability, especially for behavioral experiments that are carried out at “room temperature” and that could be influenced by temperature. To address this source of uncontrolled variability, we worked with the University to custom engineer the heating, ventilation, and air conditioning within our behavioral room so that the temperature could be precisely maintained at a set temperature (we chose 23°C) without fluctuations over the course of the day and seasons (Figure 1B,C). This temperature controlled room, and a cold room set at 4°C, were used for all subsequent behavioral studies.

**Figure 1.**
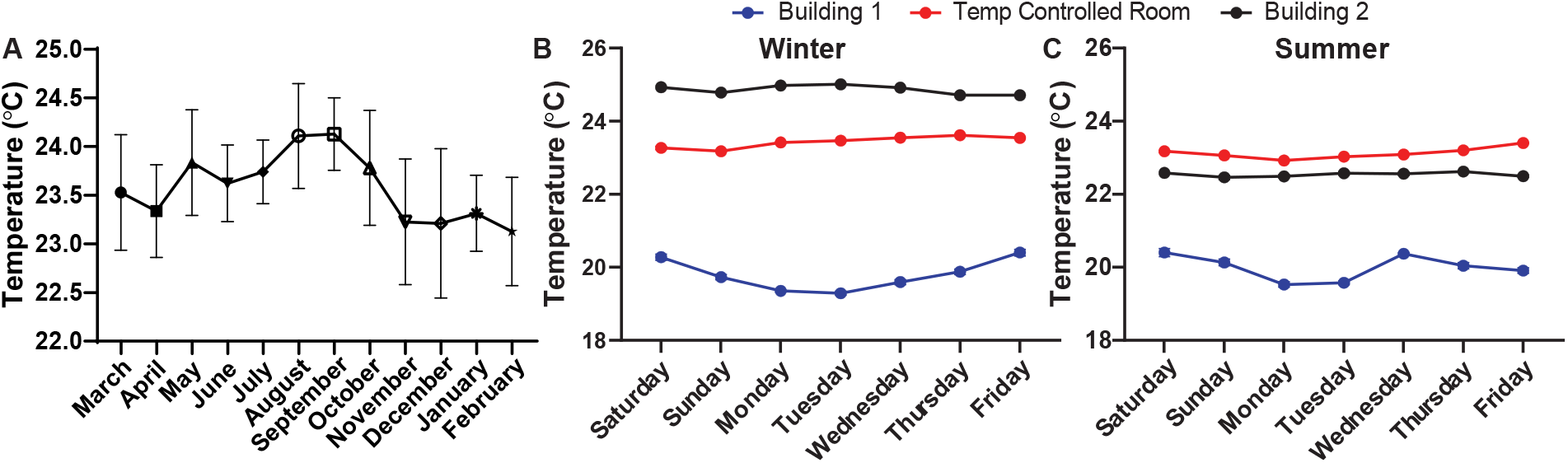
Room temperature fluctuates in lab space throughout the year if not purposefully controlled. (A) The average temperature in a semi-temperature controlled room for each month of one year. Temperature during the (B) winter and (C) summer measured every 20 minutes for one week within lab space, located in two different buildings and in a room that was specifically engineered to maintain temperature with minimal fluctuations over hours, days, weeks, and years. Temperature monitored with a La Crosse Technologies weather station.

### TRPM8 agonist impacts CBT and reduces open field behavior in WT mice

TRPM8 is a principal sensor of cold temperatures in mammalian primary sensory neurons (12, 13). To explore the impact of TRPM8 stimulation on open field behavior, we administered (i.p.) 50 mg/kg bw icilin, a TRPM8 agonist, or DMSO control to WT mice. Icilin led to a significant increase in CBT beginning at 90 minutes post injection (Figure 2A). To assess the impact of increased CBT on open field behavior, we administered 50 mg/kg bw icilin, waited 10 minutes, and then measured distance traveled and time spent in the center (Figure 2B-C). Icilin administration significantly reduced activity in the open field, suggesting that an increase in CBT led to a reduction in open field behavior.

**Figure 2.**
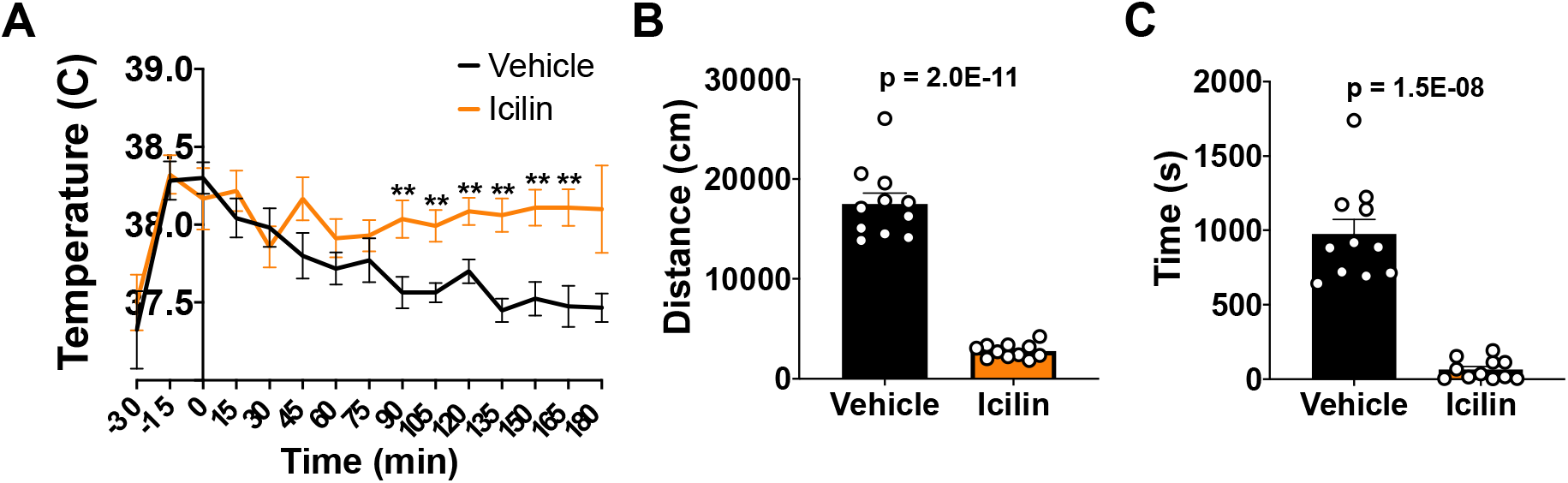
Effect of icilin on CBT and open field behavior in WT mice. CBT was measured every 15 minutes following 50 mg/kg bw i.p. administration of icilin or vehicle (A). Distance traveled (B) and time spend in the center (C) in the open field at 23°C was measured 10 minutes post-icilin administration for one hour. Data represent means ± SEM. n=11-14 mice.

### Cooling environment to 4°C reduces open field behavior in WT but not Trpm8^-/-^ mice

Stimulation of TRPM8 channels with icilin impacts the behavior of WT mice in the open field (Figure 2). Thus, we hypothesized that cold stimulation of TRPM8 channels would similarly impact open field behavior and that mice lacking TRPM8 channels would resist the effect of cold stimulation on open field behavior. We found that WT and *Trpm8*^-/-^ mice display similar distance traveled and time spent in the center when tested at 23°C (Figure 3A-B). When mice were tested at 4°C, an effect of genotype was revealed, in which the WT mice display reduced distance traveled and center time compared to *Trpm8*^-/-^ mice.

**Figure 3.**
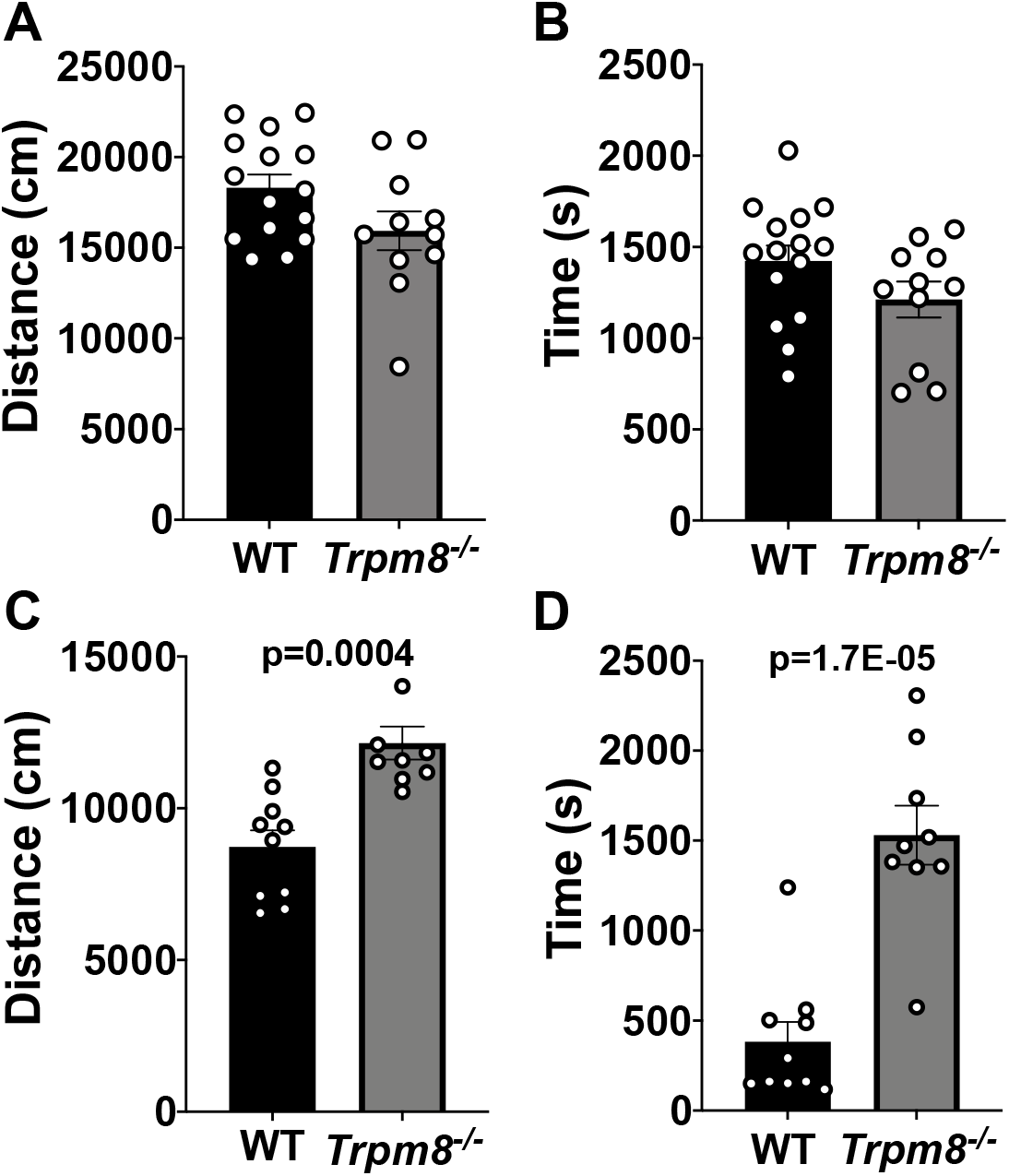
Effect of temperature on open field behavior of WT and *Trpm8*^-/-^ mice. Distance traveled (A) and center time (B) were assessed in WT and *Trpm8*^-/-^ mice at room temperature (23°C). Open field behavior was reassessed in the cold room, at 4°C (C-D). Data represent means ± SEM. n=8-15 mice.

### M8-B antagonism of TRPM8 channels increases open field behavior in WT mice

We next use a TRPM8 antagonist (M8-B) to block cold-induced stimulation of TRPM8 channels in WT mice. Administration (i.p.) of M8-B at 12 mg/kg bw decreased the CBT of WT but not *Trpm8*^-/-^ mice at >1 h post injection at room temperature (Figure 4A). Thus, mice were placed in the open field chamber 1 h following M8-B administration. We found that TRPM8 antagonist administration partially recovered the reduction in open field behavior at 4°C in WT mice (Figure 4B-C, Figure 6A-B). These data suggest that environmental temperature sensation influences open field behavior in mice.

**Figure 4.**
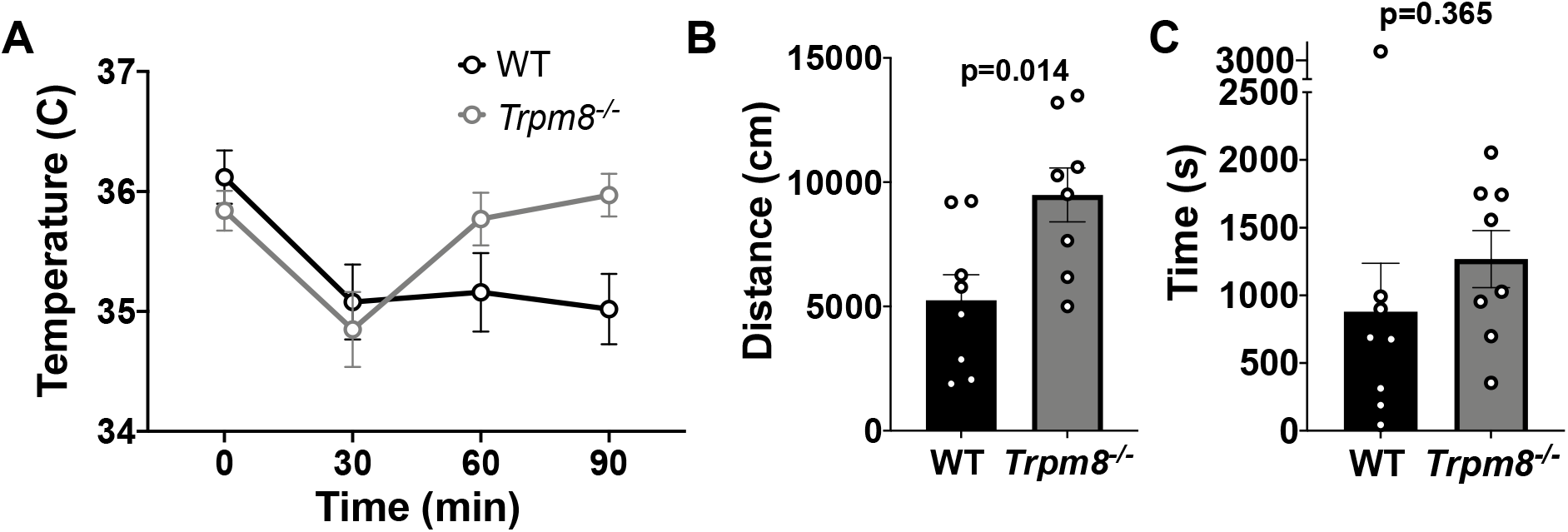
Open field behavior following TRPM8 antagonist administration (M8-B). Effect of 12 mg/kg bw M8-B on WT and *Trpm8*^-/-^ mice CBT. Total distance traveled (A) and center time (B) at 4°C 1 h following 12 mg/kg bw M8-B administration. Data represent means ± SEM. n=8-10 mice.

**Figure 5.**
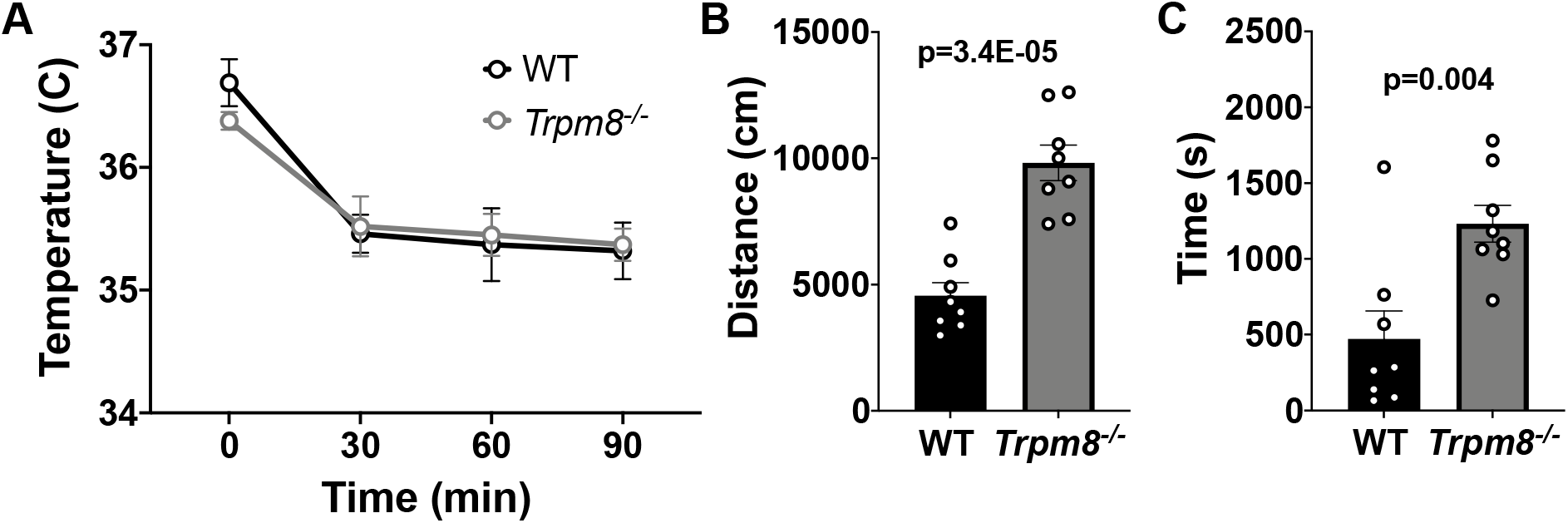
Effect of diazepam on CBT and open field behavior. CBT measured at 30, 60 and 90 minutes post 2 mg/kg bw diazepam in WT and *Trpm8*^-/-^ mice (A). Total distance traveled (B) and time spent in center (C) at 4°C, 30 minutes post 2 mg/kg bw diazepam administration. Data represent means ± SEM. n=8-10 mice.

**Figure 6.**
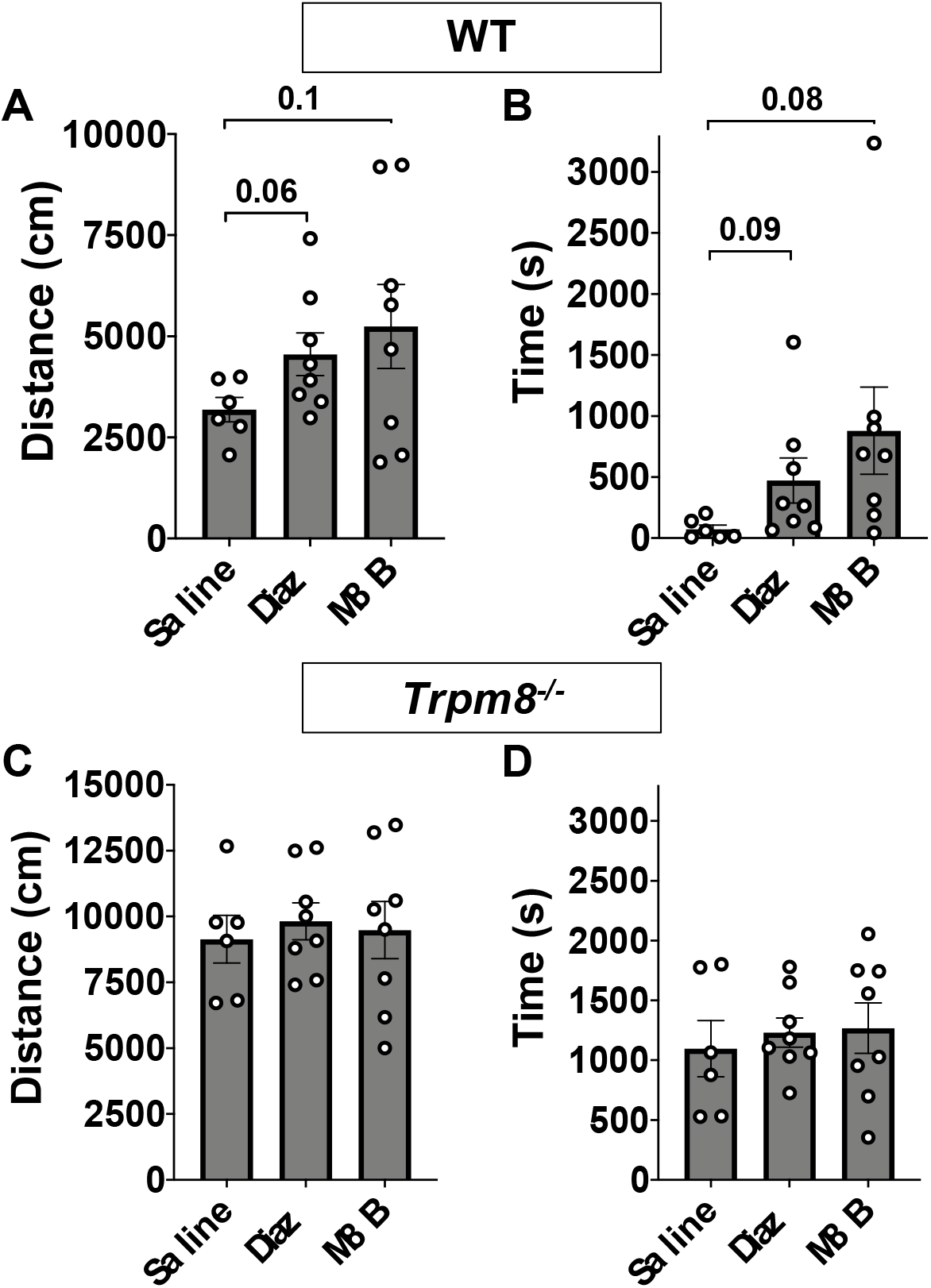
Comparison of open field behavior at 4°C in WT and *Trpm8*^-/-^ mice following saline, diazepam, or M8-B administration. Distance traveled (A) and center time (B) in WT mice. Distance traveled (C) and center time (D) in *Trpm8*^-/-^ mice. Data represent means ± SEM. n=6-8 mice.

### The anxiolytic diazepam reduces CBT in WT and Trpm8^-/-^ mice

Benzodiazepines, such as diazepam, are commonly prescribed to reduce anxiety in humans. Diazepam functions to increase gamma-aminobutryic acid (GABA) in the brain and is used to treat anxiety. To investigate whether the anxiolytic effect of diazepam was associated with a reduction in CBT in mice, WT and *Trpm8*^-/-^ mice were administered 2 mg/kg bw of the drug at room temperature. Diazepam exposure led to a reduction in CBT at 30 – 90 minutes post injection (Figure 5A). This reduction in CBT was associated with a near-significant increase in distance traveled and center time displayed by WT but not *Trpm8*^-/-^ mice, when tested at 4°C (Figure 5B-C, Figure 6).

## Discussion

In this study we compared the behavioral effects of the anxiolytic drug diazepam, with the effects of other drugs that alter cool temperature sensation and CBT, including icilin and M8-B in WT and *Trpm8*^-/-^ mice. We observed CBT and open field behavioral effects of icilin and M8-B in WT mice, but not *Trpm8*^-/-^ mice.

The effect of drugs on core body temperature may be mediated by acting on any component of the thermoregulatory system. These components include heat production, heat conservation, and thermosensing-related pathways within the nervous system that coordinate thermoregulation (8). Clark et al. present a thorough study of drug-induced changes in body temperature and provide a source of information on interactions between certain drugs and the thermoregulatory system. The data present an extensive review of the magnitude of body temperature changes induced by psychoactive compounds while taking into account the species, administration route, dose, and environmental temperature differences. Considering the effects of drug-induced changes on body temperature and the impact of CBT on behavior, studies using rodent models of psychological disorders should consider potential alterations to the perception of environmental temperatures.

The spontaneous behavior of animal models is often used to evaluate the efficacy of drugs used to treat neuropsychiatric disorders. One main concern with animal models is the lack of standardization between laboratories, which can lead to results that are not reproducible. We suggest that stricter testing protocols include assessment of room temperature and control for drug-induced alterations to CBT.

By providing greater understanding of the relationship between body temperature and behavior in mice, our data highlight the importance of assessing CBT, environmental temperature, and drug-induced changes to thermoregulation. Thus, consideration of ambient and CBT is a straightforward approach to enhance rigor and reproducibility in studies of neuropsychiatric disorders.

